# Evaluation Of The Quercetin Semisynthetic Derivatives Interaction With Breast Cancer Resistance Protein

**DOI:** 10.1101/2020.09.12.292821

**Authors:** A.E. Manukyan, A.A. Hovhannisyan

## Abstract

Breast cancer resistance protein (BCRP, ABCG2) is one of the ATP binding cassette (ABC) transporter proteins which involved in multi-drug resistance of cancer therapy. BCRP is expressed in various types of human cancer and inhibiting can be a solution to overcome multidrug resistance. Significant effort has been devoted to develop treatment strategies to overcome BCRP-mediated resistance. In addition, BCRP is expressed also in many normal tissues and plays an important role in drug absorption, distribution, and elimination. In order to design an effective cancer treatment strategy, a lot of flavones and their derivatives are used as anticancer drug compounds. One of the most promising flavonols is quercetin and quercetin semisynthetic derivatives which can inhibit the expression of BCRP.

**Highlights:**

- Virtual screening using multiple docking program for consensus predictions.
- Virtual screening of the quercetin derivatives as potential inhibitors of BCRP.
- Selected derivatives bind to the main amino acids of proteins.
- Selected derivatives meet the criteria necessary for their consideration as drugs.

## 1. INTRODUCTION

Multidrug resistance (MDR) is acquired cross resistance for cancer cells to a variety of structurally and chemically diverse chemotherapeutic agents. MDR has been considered as a major barrier for successful cancer treatment. Although many mechanisms of MDR have been identified, drug efflux mediated by the ATP binding cassette (ABC) transporter proteins represents the most common and best characterized one [6]. Among the 49 ABC transporters identified, P-glycoprotein (P-gp, ABCB1), Multi-drug resistance protein 1 (MRP1, ABCC1), and breast cancer resistance protein (BCRP, ABCG2) account for most of the observed efflux transporter-mediated MDR in cancer cells [6, 7]. Various methods and approaches are used in anticancer therapy, however, the results are not very encouraging. Currently, compounds of plant origin are considered in anticancer therapy, which possess selective cytotoxic, cytostatic, anti-inflammatory, immunomodulatory and other properties with low side effects. Nevertheless, there is an actual problem for the development of new anticancer drugs [15]. Nowadays, research, development and use of secondary metabolites have been rapidly developing [1]. To design an effective cancer treatment strategy, it is essential to understand the interactions of natural molecules with their targets. Among the various natural substances, flavonoids belong to one of the most promising groups which have been utilized for cancer treatment [2, 3]. Quercetin (C15H10O7) is a lipophilic bioactive flavonol. [3]. It can cross the cellular membranes and trigger many various intracellular pathways involved in chemoprevention. Nowadays semisynthetic compounds which based on plant active compounds as quercetin are considered as potential inhibitors for BCRP.

Exploring the mechanistic insight of such bioactive compounds will help us for further stimulate the scientific community to design novel anticancer strategies shortly.

The current paper focuses on the quercetin and quercetin semisynthetic derivatives possible inhibition of BCRP. Selected derivatives meet the criteria necessary for their consideration as drugs and can serve as a basis for conducting further in vitro/ in vivo experiments. It could be used for the development of modern anticancer therapy.

## BCRP STRUCTURE AND ACTIVITY

According to the Human Gene Nomenclature Committee, BCRP is classified as the second member of subfamily G of the ABC transporter superfamily, and hence is named as ABCG2. Most ABC transporters are characterized by two ATP-binding domains. Efflux transporters in cancer resistance and two transmembrane domains [6]. BCRP consists of 655 amino acids and hydropathy analysis predicted that BCRP contains only one ATP-binding domain (amino acid residues 1–396) followed by one transmembrane domain (amino acid residues 397–655), and therefore is considered as a half transporter [6]. As a functional ABC transporter requires two ATP-binding domains and two transmembrane domains, which form a central substrate translocation pathway, so BCRP needs at least to dimerize in order to be functional [22, 23]. Although the mechanism of BCRP dimer/oligomer formation remains unclear, several studies suggested that it may involve intermolecular disulfide bonds formed between extracellular Cys residues, among which Cys603 appears to be an important residue [29, 30]. The transmembrane domain of BCRP contains six transmembrane α-helices, connected by two intracellular and three extracellular loops. It is generally believed that ABC transporters, including BCRP, have multiple drug binding sites in the large pocket formed by those transmembrane α-helices [15]. BCRP expression can be regulated at the transcriptional level, although the regulation of the ABCG2 gene is complex. In addition, BCRP expression has been found to be induced by the peroxisome proliferator-activated receptor gamma [15].

### BCRP and molecular targeted anticancer therapy

Most actual therapy for BCRP is molecular targeted anticancer therapy, which represents entirely new classes of more target specific anticancer drugs. Unlike conventional chemotherapy that can cause non discriminating damage to both normal and cancer cells, molecular targeted therapy attacks cancer-specific targets and therefore has a more favorable safety profile [91, 92].

According to a wide range of experiments, phytochemicals including flavones, as well as flavonoids possess considerable anti-cancer features, which can be employed against different kinds of cancers [6]. In this regard, has been reported pharmacologic activities for quercetin, including anti-oxidant, anti-inflammation, as well as anti-proliferation [6]. Many studies have evaluated the effect of flavonoids as potent BCRP inhibitors [9, 12].

### Quercetin

Quercetin (2-(3,4-dihy-droxyphenyl)-3,5,7-trihydroxy4H-chromen4-one) includes two benzene rings named A and B, and joined through a 3-carbone heterocyclic pyrone one [9] (Fig. 1). Since two antioxidant pharmacophores are present in the structure of quercetin, it can largely remove free radicals and join to transitional metal ions. Moreover, catechol along with the OH group presenting at the position C3 in the structure of quercetin is an ideal arrangement to scavenge free radicals [9]. This agent is a pentalhydroxyl-flavonol consisting of 5 hydroxyl groups on the flavonol structure at 3, 30, 40 5, and 7 position carbons. Replacement of various functional groups leads to different biochemical as well as pharmacologic properties of quercetin [11]. Significant number of studies focused on anti-cancer properties of this bioactive compound. Several pathways have been identified which are affected by quercetin in different cancer [11, 12]. Based on available evidences, quercetin can inhibit a broad range of cancer such as breast, lung, nasopharyngeal, kidney, colorectal, prostate, pancreatic, as well as ovarian cancers. Quercetin is not harmful for healthy cells, while it can impose cytotoxic effects on cancer cells through several mechanisms, making it and it’s derivatives as good candidates to treat cancer by inhibiting BCRP [].

**Fig.1.**
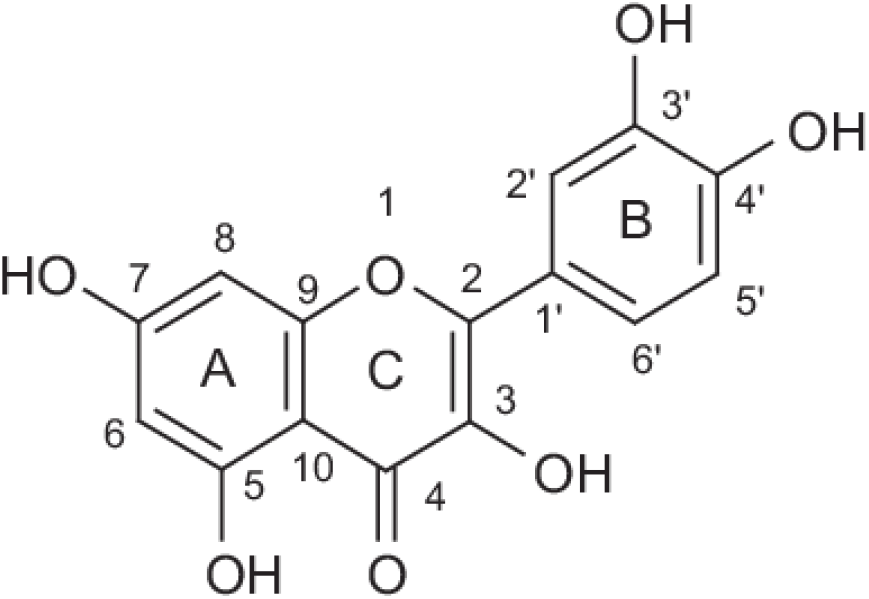
Structure of the Quercetin.

### Quercetin derivatives

As material for *in silico* analysis we took studies of Massi *et al.* which is about developments in the synthesis and anticancer-related activities of quercetin derivatives reported from 2012 to 2016 [11] (fig 2).

**Fig.2.**
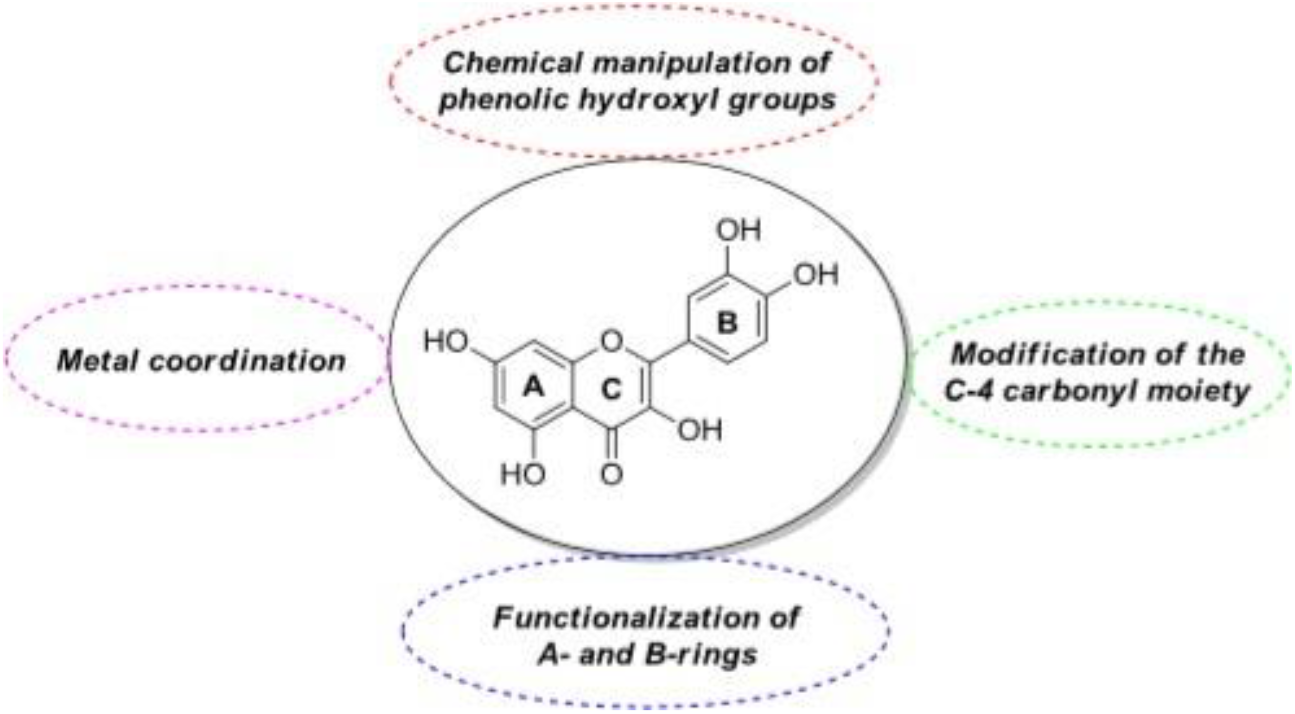
General manipulations with quercetin structure.

## 2. MATERIALS AND METHODS

### 2.1. Preparation of the protein structure and identification of binding sites

Crystallographic data on the structure of proteins belonging to *Homo sapiens* were taken from PDB RCSB (PDB: 4FYQ) [2]. Linking sites verified using MetaPocket server. As a result, 2 major binding pockets were identified in protein: in the nucleotide-binding domain (NBD).

### 2.2. Preparation of the chemical compounds library

Quercetin is a secondary metabolite that exhibits a wide range of pharmacological activities: anti-inflammatory, antiviral, antineoplastic, antibacterial, etc. To create a library of chemical compounds, the structure of quercetin was obtained from the Chemical compounds database-PubChem, 73 compounds which already synthesized as derivatives [11], were constructed using MarvinSketch [11, 12]. Compound structures were tested based on the five Lipinski rule. For the preparation and the optimization of the ligands the conformational mobility (determination of the degrees of freedom) and charges for all atoms were calculated, polar hydrogen atoms were inserted. The structures were processed out using the software package OpenBabel v2.4.0 [4].

### 2.3. Molecular docking

Molecular docking was performed using the AutoDock Vina and LeDock software package [16]. This two software packages were chosen based on their accuracy [17].

The whole protein conformational space was searched, using Grid box dimensions 94×102×90 Å for Autodock Vina. A total of 40 docking trials were performed. Following exhaustiveness values were tested in this study: 8, 16, 32, 64, 128, 256, 512, 1024, 2048 and 4096. The number of interaction sites does not change in the interval using exhaustiveness from 1024 to 4096. Exhaustiveness value of 1024 was chosen as it provides good results, good speed and thorough sampling of the docked configurations. Exhaustiveness value was increased to a value of 1024, and a maximum number of binding modes to generate set to 20. After that 50 independent docking calculations were carried out with random initial seeds. The radius was to 60 Å to cover the whole protein surface. The box size of LeDock was also set to cover the whole protein surface with the following values: xmin = −10, xmax = 11; ymin = −20, ymax = 21; zmin = −42, zmax = 43.

AutoDock Tools [16], was used for the configuration of the simulations. Using this program, two docking boxes were selected, whose sizes did not exceed 28,000 Å3. The binding affinity ΔG was used, which is calculated on the basis of the total energy of intermolecular forces, including Van der Waals forces, electrostatic, hydrophobic interactions and hydrogen bonds. Low values of ΔG correspond to strong, while high values correspond to weak ligand binding [16].

### 2.4. Similarity analysis of drugs

Calculation of physicochemical descriptors, as well as the prediction of ADME properties: Absorption, Distribution, Metabolism, Excretion and Toxicity, were determined using two different programs: SwissADME [5] and PASS online [18], which are used for drug discovery. The toxicity prediction on cell lines was also carried out using PASS online [28], which estimates the predicted types of activity with the estimated probability for each type of activity “to be active” Pa and “to be inactive” Pi, which vary from zero to one.

## 3. RESULT and DISCUSSION

After analysis from 73 compounds were selected 10 compounds that received high marks according to the results of docking analysis (Fig. 5). Unfortunately, molecular docking does not provide an unambiguous solution to the search for drug compounds, however, comparing the studies already completed and tested *in vitro*, the results are encouraging. In the studies of Yuan et al, it was shown that these ten compounds can be inhibitors of BCRP [11].

### Binding constants and the number of binding sites

An important feature was that the replacement of the quercetin structure at the 3’, 4’-O-Met positions, as well as the addition of the 5-OH group during in vivo studies became necessary - they increased the inhibitory properties of the new derivatives.

Shi and his colleagues studied O-methyl compounds, which can inhibit the growth of cancer cells, as shown in their studies [13]. These studies have shown that selective masking of the hydroxyl groups of quercetin is crucial in determining antiproliferative activity. In general, it was possible to maintain an inhibitory effect against cancer cell lines by methylation at the 4 / OH and / or 7-OH positions, while substitution of the 3’- and 4’-O-Me groups increased the activity of the derivatives.

As a result of in silico studies, it was found that the compounds binding to BCRP form bonds with the Arg482, Pro485, Thr402, Cys603, Asn629, Asn387 NBD site, which may indicate the possibility of competitive interaction of these compounds.

It should be noted that as a result of docking analysis, other compounds were found that showed high affinity and which can strongly bind with the formation of hydrogen bonds or due to van der Waals forces, electrostatic interactions with amino acids, but since in vitro studies during Since the studies refuted the activity of these compounds, the latter are not given in this article. Arg482 exhibits transport activity as well as substrate specific activity; binding to this amino acid results in protein inhibition. Pro485 is an important amino acid, the inhibition of which leads to the inhibition of efflux activity, mutations or interactions close the protein channel, as well as substrate-specific activity. The Thr402 amino acid is responsible for the ATPase activity of the protein, binding to which leads to the decomposition of the protein complex. Cys603 is involved in the formation of the dimeric structure of the protein; binding to the latter leads to the fact that the protein cannot dimerize, thereby not forming an active complex. Asn387 and Asn629 can be involved in substrate interaction.

### Analysis of the similarity of in silico research methods with in vitro methods

According to the calculated parameters, the selected 10 compounds had potential properties for use as a medicine in biological systems. SwissADME [18] predicted a high probability of adsorption by the gastrointestinal tract (HIA +) and penetration through the blood-brain barrier (BBB +) except for derivatives 15 and 73 for which penetration through the blood-brain barrier (BBB−) was not predicted. According to the predicted PASS results, the online compounds are not toxic, the compounds meet the criteria for drug similarity. In silico results display in vitro results. Thus, these compounds deserve further in vivo.

## CONCLUSION

There are a large number of flavones and their derivatives, which can be considered as potential antitumor drug compounds that can inhibit the activity of BCRP protein. Compounds that bind to amino acids that play an important role in protein expression have been identified, which is of interest for further study of these compounds. In the course of the work done, it was found that *in silico* methods can be obtained, the results are consistent with the results obtained in vitro methods. The selected compounds meet the criteria necessary for their consideration as medicines, and can serve as the basis for further *in vivo* experiments.

**Table 2.** Results for ABCG2.

| *Ligand-protein binding results* |  |  |
| --- | --- | --- |
| Derivatives* | Gibbs free energy<br>(kcal/m) | Binding constant (10<sup>6</sup>) |
| 4 | -8.67 | 21,1 |
| 5 | -9.32 | 62,8 |
| 8 | -8.72 | 22,9 |
| 15 | -9.32 | 57,7 |
| 21 | -9.45 | 71,8 |
| 26 | -8,95 | 62,8 |
| 29 | -9.22 | 53,1 |
| 51 | -9.33 | 60,7 |
| 53 | -9.12 | 43,4 |
| 73 | -8.55 | 17,2 |

